# The delayed bloodstream clearance of *Plasmodium falciparum* parasites after M5717 treatment is attributable to the inability to modify their red blood cell hosts

**DOI:** 10.1101/2023.04.24.538013

**Authors:** Molly Parkyn Schneider, Oliver Looker, Maria Rebelo, David K. Khoury, Matthew W.A. Dixon, Claude Oeuvray, Brendan S. Crabb, James McCarthy, Paul R. Gilson

**Author notes:** Dr. Maria Rebelo. Instituto de Medicina Molecular Joao Lobo Antunes, Lisboa, Portugal.

## Abstract

M5717 is a promising antimalarial drug under development thatacts against multiple stages of the life cycle of *Plasmodium* parasites by inhibiting the translation elongation factor 2 (*Pf*eEF2), thereby preventing protein synthesis. The parasite clearance profile after drug treatment in preclinical studies in mice, and clinical trials in humans showed a notable delayed clearance phenotype whereby parasite infected red blood cells (iRBCs) persisted in the bloodstream for a significant period before eventual clearance. In a normal *P. falciparum* infection iRBCs sequester in the deep circulation by cytoadherence, allowing them to avoid surveillance and clearance in the spleen. In this work we show that M5717 treatment renders iRBCs invisible to normal splenic clearance mechanisms. We found that M5717 blocks parasite modification of their host red blood cells (RBCs) by preventing synthesis of new exported proteins, rather than by directly blocking the export of these proteins into the RBC compartment. Using *in vitro* models, we demonstrated that M5717 treated ring/trophozoite stage iRBCs became less rigid, and cytoadhered less well compared to untreated iRBCs. This indicates that *in vivo* persistence of M5717 treated iRBCs in the blood stream is likely due to reduced cytoadherence and splenic clearance.

## 1 Introduction

Malaria remains a major global health burden that has recently worsened due to the impact of the SARS-CoV-2 pandemic, with an estimated 247 million cases of malaria and 619,000 deaths in 2021 (World Health Organization, 2022). This represents a 10% increase in malaria associated deaths from previous years. Compounding the challenge of malaria elimination is increasing resistance to frontline artemisinin derivatives. Resistance alleles first appeared in the Greater Mekong subregion (Dondorp et al., 2009), but have more recently been observed in Africa, South America and Papua New Guinea (Bergmann et al., 2021;Lautu-Gumal et al., 2021;Yoshida et al., 2021;Ward et al., 2022). Artemisinin derivatives have rapid killing action but a short half-life in the body, so are partnered with long-lasting antimalarials. In addition to the spread of resistance to artemisinin drugs, resistance to partner drugs is also increasing (Marapana and Cowman, 2020). It is therefore essential to develop new antimalarials to replace both the artemisinin derivatives and their partner drugs. New drugs should ideally act upon more than one lifecycle stage (namely the liver and transmission stages), have a novel mechanism of action to reduce cross-resistance, and be affordable (Burrows et al., 2017).

M5717 (previously known as DDD107498) is a promising new antimalarial. It has nanomolar potency against the asexual blood stages of *Plasmodium falciparum* and satisfies many partner drug criteria, with an ability to kill gametocytes and block liver stage infections in mouse models of malaria (Baragana et al., 2015;Baragana et al., 2016). The molecular target of M5717 is the parasite’s translation elongation factor 2 (*Pf*eEF2), disruption of which inhibits protein synthesis (Baragana et al., 2015). This, along with its long half-life in humans (146–193 hr) (McCarthy et al., 2021), makes M5717 an appealing drug for a single-dose cure of malaria. One potential shortcoming of M5717 however, is that the minimum inoculum for resistance (MIR) for cultured *P. falciparum* is 5 x 10^6^ parasites, which means that its refractiveness to resistance is only modest (Duffey et al., 2021).

In studies undertaken in a severe combined immunodeficient (SCID) mouse model of *P. falciparum* malaria, M5717 was slow acting, but performed well when partnered with pyronaridine, a faster acting drug which inhibits haem metabolism (Rottmann et al., 2020).

In a recent phase 1a/b trial in human volunteers, the blood stage activity of M5717 was tested in a single dose regime in a volunteer infection study (McCarthy et al., 2021). The drug was very well tolerated up to 1250 mg, above which moderate and transient neurotoxicity was reported, namely blurred vision and oral numbness. Interestingly, the parasite clearance profile was biphasic with an initial slow clearance lasting 35-55 hours, followed by a rapid clearance phase. Recrudescence occurred in some participants at the lower 150 mg and 400 mg doses, where *in vivo* selection for genetic mutations in *Pf*eEF2 was also observed (McCarthy et al., 2021).

*P. falciparum* is predicted to export hundreds of proteins into its RBC compartment, many of which facilitate avoidance of clearance by the innate and adaptive human immune system (Hiller et al., 2004;Marti et al., 2004;Sargeant et al., 2006;Boddey et al., 2013;Jonsdottir et al., 2021). These exported proteins are first delivered by the vesicular trafficking pathway to the parasitophorous vacuole (PV) encasing the intraerythrocytic parasite. Proteins are subsequently translocated into the RBC compartment via the Plasmodium Translocon of Exported proteins (PTEX) (de Koning-Ward et al., 2009;Beck et al., 2014;Elsworth et al., 2014;Ho et al., 2018).

Here we aimed to determine why M5717 treated parasites persist in circulation without being cleared for 48 – 60 hours after treatment. We hypothesised that M5717 treatment results in defects in the trafficking and/or export of proteins responsible for RBC membrane skeleton remodelling and sequestration. In the absence of this remodelling the biophysical properties of the iRBC are altered, allowing for partially remodelled non-sequestering cells to remain in the circulation and avoid splenic clearance.

## 2 Materials and Methods

### 2.1 Parasite Culture and Strains

*Plasmodium falciparum* CS2, wild-type 3D7 and transgenic 3D7 parasites were continuously cultured at 4% haematocrit (HCT, Australian Red Cross Lifeblood) in RPMI medium (Sigma Aldrich) supplemented with 0.2% NaHCO_3_ (Thermo Scientific), 0.5% Albumax II (GIBCO), 25 mM HEPES (GIBCO), 0.37 mM hypoxanthine (Sigma Aldrich) and 31.25 µg/mL Gentamicin (GIBCO) (Complete Media). Cultures were kept at 37 °C in the presence of 1% O_2_, 5% CO_2_, 94% N_2_. The exported Hyp1-Nluc and secreted MSP1-Nluc bioluminescent proteins were expressed from plasmids transfected into 3D7 as previously described (Azevedo et al., 2014) and were maintained with 2.5 nM WR99210.

### 2.2 M5717 Preparation and Growth Assay

The compound M5717 (Merck) was dissolved in 100% sterile DMSO to a concentration of 10 mM. Stock aliquots were made to a concentration of 250 µM and diluted further to 250 nM in Complete Media. To determine the EC_50_ a 72-hour growth assay was performed on synchronised ring stage 3D7 parasites at 2% HCT and 0.3% parasitemia in a 96 well U-bottom plate. Serial dilution of M5717 by 2-fold and over 9-points, starting at the highest concentration 10 nM, was performed. Uninfected RBCs and 0.05% DMSO were used as controls. The parasites were incubated at 37 °C for 72 hours followed by a lactate dehydrogenase assay to determine parasite proliferation (Makler and Hinrichs, 1993). Samples were measured in triplicate, background signal was measured from the RBC control and subtracted from each group before normalisation to the DMSO control. The concentrations were then log transformed and Graphpad Prism used to perform a nonlinear regression (log(inhibitor) vs response = 4 parameters).

### 2.3 Ex vivo growth of *P. falciparum* parasites after M5717 administration

All reagents were purchased from Life Technologies Australia, unless stated otherwise. The *ex vivo* experiments presented here were performed as an exploratory study of malaria volunteer infection study randomised clinical trial (NCT03261401) (McCarthy et al., 2021). The study was done at Q-Pharm (Brisbane, QLD, Australia) following approval by the QIMR Berghofer Medical Research Institute Human Research Ethics Committee. All participants gave written informed consent before enrolment. The study is registered with ClinicalTrials.gov (NCT03261401). Samples and data were obtained from cohort 3 of the randomised clinical trial (McCarthy et al., 2021). Subjects were inoculated with approximately 1,800 viable *P. falciparum* 3D7 infected red blood cells. Eight days after inoculation subjects received a single oral dose of 800 mg of M5717. Parasitaemia in the volunteers was monitored by a qPCR assay targeting the *P. falciparum* 18S rRNA gene. Blood samples were collected in EDTA tubes (Becton Dickinson, Australia) before M5717 administration (t = 0 h) and at 24, 48 and 72 h post-M5717 administration from all subjects (n = 8) and cultured *ex vivo*. One participant was excluded from this analysis because parasitaemia was very low (13 parasites/mL) on day of dosing. The blood samples collected from subjects infected with *P. falciparum* and administered with M5717 were washed at 37 °C in 25 mL of RPMI 1640 medium and centrifuged at 530 g for 5–10 minutes. A volume of 250 μL of the RBC pellet was resuspended in 5 mL of RPMI 1640 medium supplemented with 10% heat inactivated pooled human serum and 0.1 mM of sodium hypoxanthine to obtain approximately 5% haematocrit. Cultures were established in 6 well-plates and maintained at 37 °C, in a 5% CO_2_, 5% O_2_ and 90% N_2_ atmosphere. A volume of 50 μL of uninfected RBCs (Australian Red Cross Blood Service), at 50% haematocrit, was added to the cultures every 6 to 8 days after the pre-M5717 culture (t = 0 h) was established. Parasitaemia in *ex vivo* cultures, defined as the percentage of infected RBCs, was monitored by flow cytometry for up to 25 days. Parasitaemia was determined by assessing the percentage of SYBR green I positive cells. For each measurement, 5 μL of the culture was stained with SYBR green I (diluted 1:1000 in PBS supplemented with 2% FBS) and incubated for 30 minutes in the dark, followed by flow cytometry analysis using a BD FACS CANTO II flow cytometer. Flow cytometry data was analysed using FlowJo software (version 10.8, Tree Star Inc., Oregon, USA). The time for each *ex vivo* culture to reach a parasitemia of 0.2% was determined by log-linear interpolation between the last time point where parasitemia was below 0.2% and the first time point where parasitemia was above 0.2%. The arithmetic mean time was calculated for all samples collected pre- and post-administration of M5717.

### 2.4 Parasite Viability Assay

Sorbitol synchronised Hyp1-Nluc parasites were diluted into a 24-well plate at 4% HCT, 2% ring stage parasitemia with Complete Media. Compounds were added to separate wells in the following concentrations: 1x EC_50_ M5717 (0.56 nM), 5x EC_50_ M5717 (2.8 nM), 10x EC_50_ M5717 (5.6 nM), 5x EC_50_ artemisinin (40 nM), 5x EC_50_ chloroquine (64.5 nM), 100 µM cycloheximide and 0.05% DMSO. A blood smear was immediately taken (0 h), fixed in methanol for 30 s and stained in 10% Giemsa for 5 min. An aliquot of resuspended culture was removed at the same time and frozen in a 96-well plate at -20 °C. Parasites were returned to 37 °C. Over the following 72 hours, another blood smear and aliquot were taken at each 24 hour time point. Results were determined by counting 1000 cells for each treatment and time point, separating parasites based on their appearance into either specific blood-stage or if abnormal and/or dead. Images were taken with a Leica IC50 HD camera attached to a Leica DM750 microscope. Nanoluciferase bioluminescence was measured from the Hyp1-Nluc parasites by lysing the parasite culture 1:5 with 1x Nano-Glo Lysis Buffer (Promega, USA), and the addition of 1:1000 Nano-Glo (Promega). The bioluminescent signal was measured using a CLARIOstar luminometer (BMG Labtech) at 3000 gain for 1 s/well.

### 2.5 Counterscreen and Translation Inhibition Assay

A counterscreen was performed on the Nluc expressing Hyp1-Nluc parasite line. Parasites were treated with 25 nM ML10 (LifeArc) to prevent egress and then separated on a 67% Percoll density gradient in RPMI before resuspension in Complete Media with fresh RBCs. Cultures were shaken at 50 rpm at 37 °C for 4 hours to allow for invasion, followed by sorbitol lysis and growth at 37 °C for 20 hpi (±2 h). In a 24-well plate, the parasite culture was diluted to 2% HCT, 1% parasitemia and lysed in 1x Nano-Glo Lysis Buffer (Promega, USA), before being added to a white 96-well plate in triplicate. The following compounds were added to separate wells: 1x EC_50_ M5717 (0.56 nM), 5x EC_50_ M5717 (2.8 nM), 10x EC_50_ M5717 (5.6 nM), 10 µM DH11 (MMV665972) and 0.1% DMSO. Lysates were incubated for 10 min at 37 °C. After the addition of 1x Nano-Glo Lysis Buffer and 1:1000 Nano-Glo substrate (Promega, USA), relative light units (RLU) were measured by luminescence using a CLARIOstar luminometer (BMG Labtech) at 3000 gain for 1 s/well. Data was analysed and graphed using Graphpad Prism.

The translation assay was then performed in a similar manner, where parasite cultures were treated with the compounds 1x EC_50_ M5717, 5x EC_50_ M5717, 10x EC_50_ M5717, 100 µM cycloheximide, 5x EC_50_ artemisinin (40 nM), 5x EC_50_ chloroquine (64.5 nM), and 0.1% DMSO. Parasites were then grown at 37 °C and samples taken at time points 0 h, 2 h and 5 h of treatment. Aliquots of 5 µL parasite cultures were transferred to a white 96-well plate and mixed with 45 µL 1x Nano-Glo Lysis Buffer and 1:1000 Nano-Glo substrate. RLU was measured by a CLARIOstar luminometer (BMG Labtech) at 3000 gain for 1 s/well. Data was analysed and graphed using Graphpad Prism.

### 2.6. Export Assay

Hyp1-Nluc parasites were synchronised as described for the counterscreen. Five separate cultures of Hyp1-Nluc, and one 3D7 culture were prepared at 2% HCT and 1% parasitemia in Complete Media. The Nluc-expressing parasite cultures were each treated with one of the following: 1x EC_50_ M5717 (0.56 nM), 5x EC_50_ M5717 (2.8 nM), 5 µg/mL Brefeldin A (BFA) or 0.1% DMSO. Parasites were then aliquoted into 4 x 96-well round-bottom plates and allowed to grow at 37 °C. Plates were successively removed from incubation and stored at 4 °C at four timepoints: 0 h, 1 h, 2.5 h or 5 h. The Nluc export assays were then performed as previously described (Charnaud et al., 2018). Graphpad Prism was used to plot the data and standard error of the mean.

### 2.7 Microfiltration

MSP1-Nluc parasites were synchronised to 15 hpi (±1h) as described in the counterscreen and diluted to 3-5% parasitemia and 2% HCT. The parasites were treated for 5 h with 5x EC_50_ M5717 (2.8 nM) or <0.001% DMSO (equivalent volume), before resuspension in RPMI. In triplicate, a syringe pump (Pump 11 Elite, Harvard Apparatus) was used to flow 800 µL of parasites through a microbead matrix (5-15 µm and 15-25 µm in diameter) at a rate of 0.8 mL/min. The flow through was collected and pelleted by centrifugation at 500 g for 4 min. Giemsa smears were prepared from the pelleted samples. To determine parasitemia 1000 cells were counted for each flow through. Data was analysed and graphed using Graphpad Prism.

### 2.8 Immunofluorescence Assay

Hyp1-Nluc parasites (18 hpi, ±2 h) or 3D7 parasites (15 hpi, ±1 hr), synchronised and treated for 5 hours as previously described in the counterscreen and translation inhibition assay, were added to 1x PBS in wells of a 24-well plate containing 12 mm round coverslips coated in 0.1% Poly-L-Lysine. The cells were allowed to settle for 30 min before fixation with 4% paraformaldehyde (Sigma Aldrich), 0.0075% glutaraldehyde (ProSciTech) in 1x PBS for 20 min. The wells were washed with 1x PBS before quenching with 0.1 M glycine, 0.02% Triton X-100 in 1x PBS for 15 min. Washing was performed twice with 0.02% Triton X-100 in 1x PBS before the cells were blocked with 1% casein, 0.02% Triton X-100 in 1x PBS for 1 hour at room temperature. Primary antibodies were prepared in blocking buffer in two combinations: rabbit anti-Nluc (1:500) (Charnaud et al., 2018) and mouse anti-EXP2 (1:500) (Bullen et al., 2012), or rabbit anti-EXP2 (1:500) (Bullen et al., 2012) and mouse anti-KAHRP (1:500) (Taylor et al., 1987) and allowed to probe overnight at 4 °C. The wells were washed with 0.02% Triton X-100 in 1x PBS after removal of the primary antibodies. Secondary antibodies were prepared in 5% goat serum in blocking buffer: goat anti-mouse 594 (AlexaFluor Invitrogen, 1:2000) and goat anti-rabbit 488 (AlexaFluor Invitrogen, 1:2000). Probing was performed for 1 h at room temperature in the dark. The cells were washed again in 0.02% Triton X-100 in 1x PBS and then mounted on glass slides with mounting media containing DAPI (Vectashield, Vector Labs, USA). Imaging was performed using a Zeiss CellObserver widefield fluorescence microscope and 63x objective. Image processing was performed using ImageJ.

### 2.9 Sorbitol Lysis Assay

Synchronous Hyp1-Nluc parasites (22 hpi, ±2 h) were diluted to 1% HCT and 1% parasitemia in three wells of a 24-well plate. Each well was treated with either 5x EC_50_ M5717 (2.8 nM), 100 µM furosemide or 0.1% DMSO for 5 h. Parasites were then resuspended and 10 µL transferred to a 96-well white plate in triplicate and a sorbitol lysis assay performed as previously described (Dickerman et al., 2016). After addition of the buffers the plate was immediately shaken for 5 seconds at 700 rpm and bioluminescence recorded at 3000 gain every 3 min for 41 min 33 s. Bioluminescence was read using a CLARIO star luminometer (BMG LabTech) and the results were calculated and graphed from the three technical replicates using Graphpad Prism.

### 2.10 Parasite binding under flow conditions

Gelatin selected CS2 parasites were synchronised to 15-20 h at 1% HCT and 3% parasitemia before 5 hours treatment with either 5x EC_50_ (2.8 nM) or <0.001% DMSO (equivalent volume). Ibidi µ-Slide 0.2 channel slides that had been treated at 37 °C overnight with either 100 µL chondroitin sulfate A (100 µg/mL; CSA; Sigma Aldrich) in 1x PBS, or only with 1x PBS, were washed with 1x PBS before blocking at room temperature with 1% BSA (w/v) in 1x PBS for 1 hour. The slides were then flushed with warm bicarbonate-free RPMI supplemented with 5% Albumax II (RPMI Alb). Immediately before binding, the treated parasite cultures were resuspended in RPMI Alb and flowed through the channels using a syringe pump at a rate of 100 µl/min for 10 min at 37 °C. The slides were then washed at the same conditions with RPMI Alb for 10 min to remove unbound RBCs. Analysis was performed by imaging 10 random fields of view (281 x 178 µm) of each condition with a 40x objective using a Zeiss Cell Observer widefield fluorescence microscope and counting bound iRBCs over three biological replicates. Counts were performed using ImageJ and results were graphed in Graphpad Prism. Ethics statement

Adapt wording from Lancet ID paper

## 3 Results

### 3.1 M5717 exposed parasites show a delayed clearance phenotype in the bloodstream

To better understand the slow clearance profile observed after M5717 administration and to investigate whether circulating parasites were rendered inviable or not by the drug, blood samples were collected from volunteers at 24, 48 and 72 hours after treatment with single oral dose of 800 mg M5717 from cohort 3 in the recent phase 1b study (McCarthy et al., 2021). Blood samples were immediately placed into *ex vivo* culture (Figure 1A), and viability of circulating parasites assessed by monitoring for parasite regrowth *ex vivo* for approximately 25 days. Parasite growth above the threshold of 0.2% parasitaemia was detected after 7.1 (95% CI: 5.8 – 8.5) days in culture in blood samples collected before M5717 administration (Figure 1B, blue traces). In samples obtained 24, 48 and 72 hours after M5717 administration, parasite growth was detected after 15.3 (95% CI: 12.6-17.9) days, 14.8 (95% CI: 11.2-18.5) days and 14.5 (95% CI: 13.1-15.9) days in culture (Figure 1B, purple, orange and red traces). This observation indicates that the development of most parasites present in circulation at 24, 48 and 72 hours after M5717 administration was arrested, and that they remained in circulation without being cleared for several days, before some began replicating. Alternatively, most parasites may have been rendered non-viable with a small number remaining viable and replicating with their exponential growth only measurable after 15 days.

**Figure 1.**
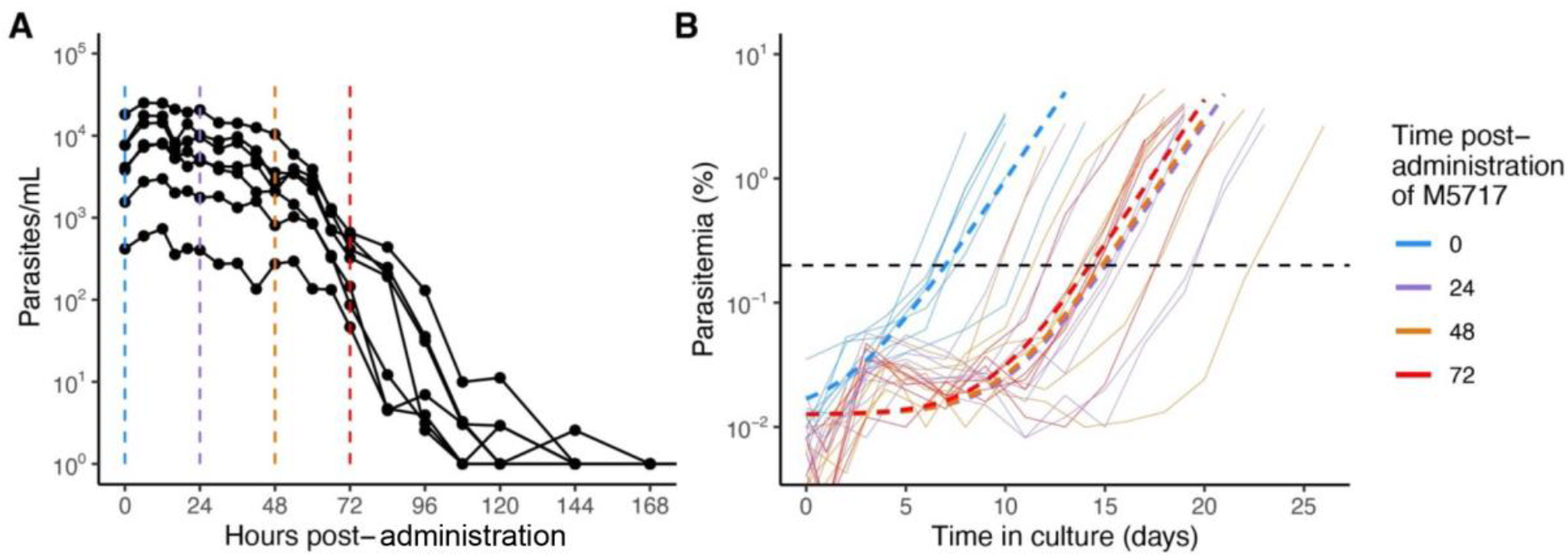
Parasitaemia after M5717 administration. **(A)** Course of parasite concentration in the volunteers determined by 18S qPCR after 800 mg M5717 administration showed a slow initial clearance profile. **(B)** Parasite levels in *ex vivo* cultures established with blood samples collected from volunteers just before (0 h in blue) and after 24 h (purple), 48 h (orange) and 72 h (red) of M5717 administration, as determined by flow cytometry (solid lines). Parasite growth above threshold of 0.2% (black dashed line) was observed after 7.1 (95% CI: 5.8 – 8.5) days in culture in samples collected before treatment, and after 15.3 (95% CI: 12.6-17.9), 14.8 (95% CI: 11.2-18.5) and 14.5 (95% CI: 13.1-15.9) days in culture in samples collected at 24, 48 and 72 h after M5717 administration. Dashed lines indicate the line of best fit obtained by fitting an exponential growth model with background constant to the individual data using non-linear mixed effects model.

### 3.2 M5717 arrests the growth of cultured *Plasmodium falciparum*

To gain insight into why blood stage parasites might persist in circulation following M5717 treatment, a 72-hour growth assay was performed with *P. falciparum* 3D7 parasites to characterise the morphological changes induced by drug treatment. The EC_50_ of M5717 was calculated to be 0.56 nM (Figure S1) and in subsequent experiments we used M5717 at concentrations that were multiples of its growth EC_50_. The effects of M5717 were compared to the more rapidly acting antimalarial compounds chloroquine (CQ) and artemisinin (Art), as well as to the protein translation inhibitor cycloheximide (CHX). The compounds were added to ring-stage transgenic 3D7 parasites expressing an exported nanoluciferase (Nluc) reporter protein (Hyp1-Nluc) and were left in for the duration of the experiment. Giemsa-stained blood smears from drug treated parasites were made daily, and the proportion of ring stage (young), trophozoite stage (mid), schizont stage (late) stage, abnormal or dead parasites was measured (Figure 2 and Figure S2). Parasites were classified as dead if they were darkly stained and shrunken (pyknotic) or were outside their RBC host cell. After 24 hours in culture, parasites treated with M5717 at 5x and 10x EC_50_ had an increased proportion of dead parasites, and transition from rings to trophozoites appeared to have stalled (Figure 2 and Figure S2). At the same time, CHX treated parasites had completely halted growth at the ring stage, whilst the more rapidly acting antimalarials Art and CQ had killed most parasites. After 48 hours parasites treated with M5717 at 10x EC_50_ were killed at a similar rate compared to Art or CQ, but at 5x EC_50_ for M5717 some parasites had progressed to trophozoite stage, indicating they may still be growing slowly. By 72 hours, growth of parasites treated with M5717 at either 5x or 10x EC_50_ had completely stalled; the remaining parasites, whilst trophozoite-like in appearance, were abnormally shaped and appeared condensed (Figure 2 and Figure S2). Parasites treated with M5717 at 1x EC_50_ appeared to grow at a similar rate compared to the DMSO control, but progression began to slow after 72 hours. Thus, compared to the fast-acting antimalarials, M5717 appeared to cause a slower form of death, in agreement with previously reported *in vitro*, animal model (Baragana et al., 2015) and clinical trial results (McCarthy et al., 2021).

**Figure 2.**
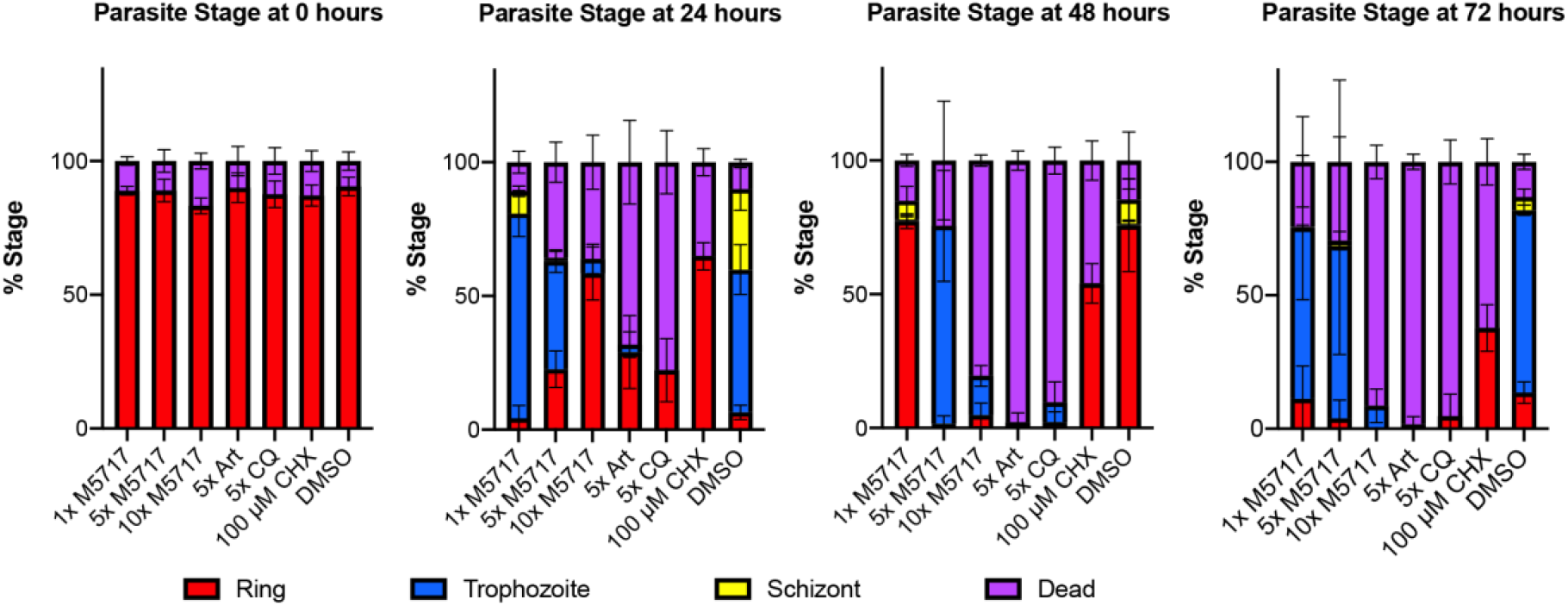
M5717 initially delays and then kills *P. falciparum* Hyp1-Nluc parasites following 24 h of treatment. Synchronous ring stage parasites were treated separately with either 1x, 5x or 10x EC_50_ of the compound M5717 (0.56 nM, 2.8 nM, 5.6 nM). Parasites were also treated with the known antimalarials artemisinin (Art, 40 nM), chloroquine (CQ, 64.5 nM), translation inhibitor cycloheximide (CHX, 100 µM), and the vehicle control DMSO (0.05%). Giemsa-stained blood smears were prepared at 0 h, 24 h, 48 h and 72 h and counted to determine the proportion of ring, trophozoite, schizont or dead/abnormal parasites. Graph represents n = 1000 cells for each treatment at each time-point. Error bars represent standard deviation of three biological replicates.

### 3.3 Optimisation of M5717 treatment for assessment of Hyp1-Nluc export

*P. falciparum* exports dozens of proteins into the RBC it infects. These significantly alter the biophysical properties of the RBC as well as cause it to adhere to the microvasculature, mediated by formation of raised structures called knobs on the RBC membrane. If M5717 blocks parasite-induced modification of the RBC membrane skeleton this could prevent parasite induced rigidity, resulting in reduced splenic clearance of the iRBCs. Likewise, if M5717 reduces cytoadherence and sequestration in the deep vasculature, this could lead to persistence of iRBCs in the bloodstream. To test if M5717 was affecting these properties, we employed parasites expressing an exported Hyp1-Nluc reporter as a marker of export. We first performed a counterscreen, which showed that M5717 does not inhibit bioluminescence of the Nluc reporter compared to the known Nluc inhibitor DH11 (MMV665972,(Dans et al., 2020)) (Figure S3A).

To quantify the effects M5717 treatment has on protein export into the iRBC compartment, we tested the compound on parasites that were still actively exporting the Hyp1-Nluc reporter. We first ensured that M5717 would not completely suppress synthesis of the reporter protein. As expression of the Hyp1-Nluc reporter begins at 18 hours post-invasion (hpi), Hyp1-Nluc parasites at 20 hpi, (± 2 h) were treated for up to 5 hours with different concentrations of M5717 or the control compounds Art, CQ and CHX. After 2 hours, there was little increase in Hyp1-Nluc activity as measured by bioluminescence (Figure S3B). After 5 hours however there was a strong increase in bioluminescence in the DMSO control, and a concentration dependent suppression of Hyp1-Nluc activity in M5717 treated parasites. This suppression was likely due to inhibition of protein translation since 10x EC_50_ M5717 produced a similar reduction as CHX. The rapid-acting antimalarial Art also caused strong suppression of Hyp1-Nluc activity, whereas CQ, which shows intermediate activity, only partially suppressed bioluminescence (Figure S3B). This suggests that M5717 would be suitable for further testing on Hyp1-Nluc parasites for 5 hours but should only be used at 1x and 5x EC_50_ to ensure sufficient new Hyp1-Nluc was synthesised for export.

### 3.4 M5717 does not directly reduce the export of parasite proteins into their RBC host cells

To measure the effects of the inhibitors on protein export, Hyp1-Nluc parasites at 20 hpi (± 2 h) were treated with M5717, CHX and DMSO as well as brefeldin A (BFA) which inhibits retrograde transport between the Golgi apparatus and the endoplasmic reticulum (Helms and Rothman, 1992). To quantify the inhibition of protein trafficking between different parasite compartments, parasitised iRBCs were collected at 0, 1, 2.5 and 5 hours after addition of M5717 and then differentially lysed. To do this, three buffers were used, Buffer A which lysed the iRBC membrane, Buffer B which lysed the iRBC and the vacuole membranes and Buffer C which lysed all membranes (Figure 3A). Following each lysis, the released Hyp1-Nluc protein hydrolysed the Nano-Glo substrate present in the surrounding buffer to produce a bioluminescence signal proportional to the amount of reporter in each lysed cell compartment. By subtracting the bioluminescent signal of Buffer B from Buffer C, the amount of Nluc signal in the PV could be ascertained (Looker et al., 2022). Hyp1-Nluc was efficiently exported into the RBC compartment in the DMSO control and displayed a time dependent retention and increase in the parasite compartment following BFA treatment (Figure 3B). In comparison, both concentrations of M5717 did not appear to specifically inhibit protein export (Figure 3B).

**Figure 3.**
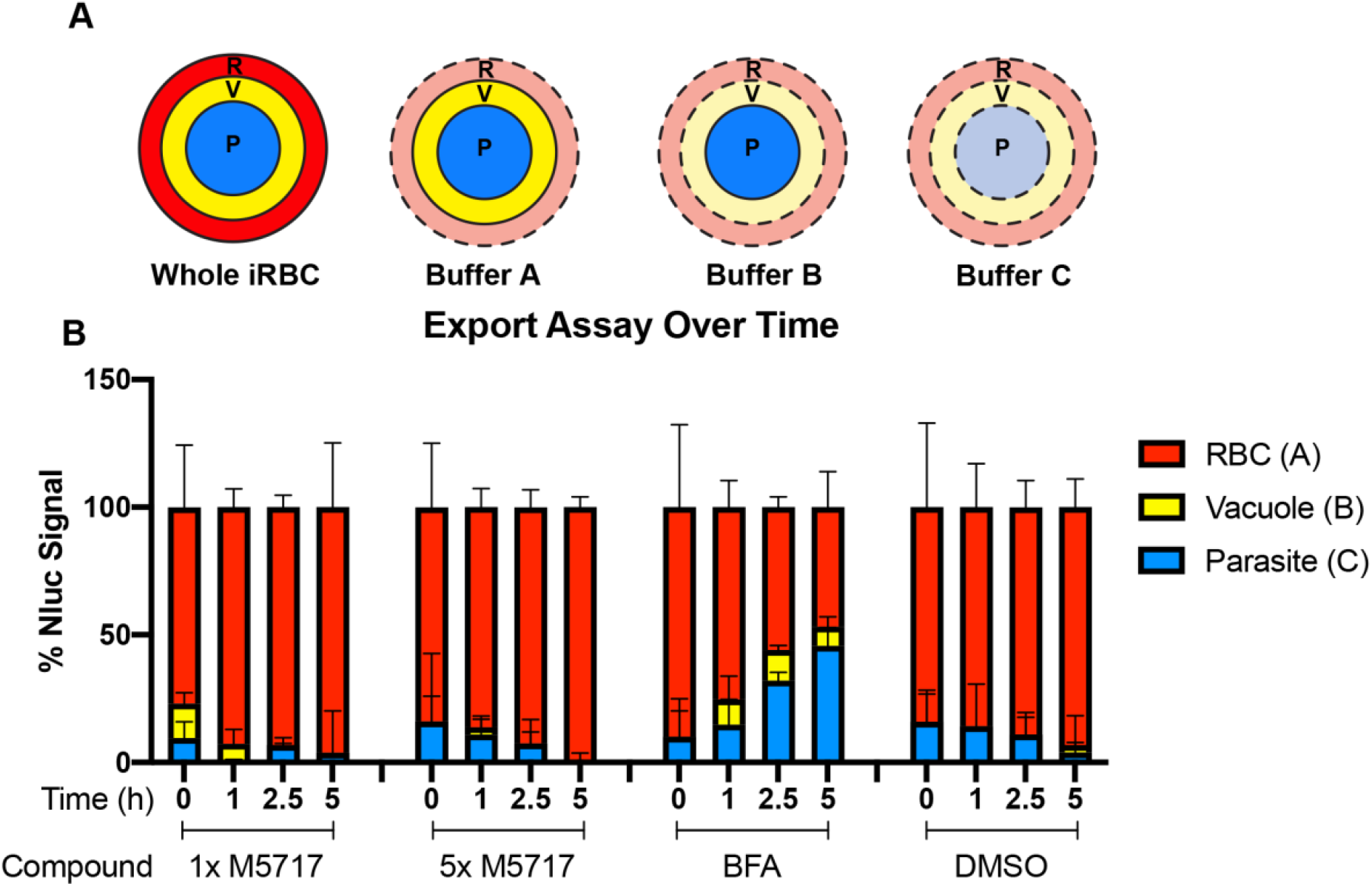
Export of nanoluciferase in *P. falciparum* 3D7 pEF-Hyp1-Nluc parasites is not specifically inhibited by the addition of M5717. **(A)** Hyp1-Nluc parasites at 20 hpi (±2 h) were treated with either 1x or 5x EC_50_ M5717 (0.56 nM or 2.8 nM), export inhibitor brefeldin A (BFA) (5 µg/mL) or vehicle control DMSO (0.1%). Samples were taken at 0, 1, 2.5 or 5 h time points and the RBC (R), RBC and vacuole (V), and all cell compartments were lysed in Buffers A, B and C, respectively. Subtraction of Buffer B signal from Buffer C indicated the nanoluciferase signal retained parasite (P) **(B)** Nanoluciferase activity was measured to determine the relative proportion of Hyp1-Nluc that was retained within each compartment for two technical replicates. Error bars represent standard error of the mean of three biological replicates.

### 3.5 Microscopic examination of M5717 treated parasites indicates exported protein expression is reduced but its delivery into the host RBC is not inhibited

Results thus far indicated that protein export is not specifically inhibited by M5717. To verify this, we further investigated protein export by fluorescence microscopy. Hyp1-Nluc Parasites at 18 hpi (± 2 h) were treated for 5 hours with 5x EC_50_ M5717 to reduce de novo Hyp1-Nluc expression. The export inhibitor BFA was used as a positive control confirming retention of Nluc signal within the parasite. Fluorescence microscopy was performed after probing parasites with antibodies for Nluc and EXP2 as a PVM marker (Figure S4). Cells were imaged randomly, and their fluorescence signal intensity was measured. The Nluc signal in the whole iRBC (Figures 4A and S4) was measured as well as the parasite signal inside the PV (Figures 4B and S4). Exported Nluc signal was calculated by subtracting the parasite signal from the whole iRBC signal (Figures 4C and S4). As anticipated, the exported signal for BFA was significantly reduced, as most protein was trapped in the parasite (Figures 4C and S4). For M5717, the parasite Nluc signal (Figure 4B) and exported Nluc signals (Figure 4C) were reduced, suggesting that expression of Hyp1-Nluc was inhibited. This was not unexpected, as the mechanism of action for M5717 is through translation inhibition. Interestingly, when treated with M5717 the ratio of Nluc exported signal to the internal parasite signal was higher than in DMSO treated cells, which contrasts with the significantly reduced signal in iRBCs treated with BFA (Figures 4D and S4). This comparison shows that the protein export mechanism has not been directly disrupted and some Hyp1-Nluc synthesised prior to M5717 treatment has still been successfully exported. Some expression of Hyp1-Nluc therefore likely begins earlier than the time-points that these experiments were initiated, which could explain why fluorescence signal persists in the RBC despite treatment with M5717. This result therefore supported the results of the nanoluciferase export assays reported above (Figure 3).

**Figure 4.**
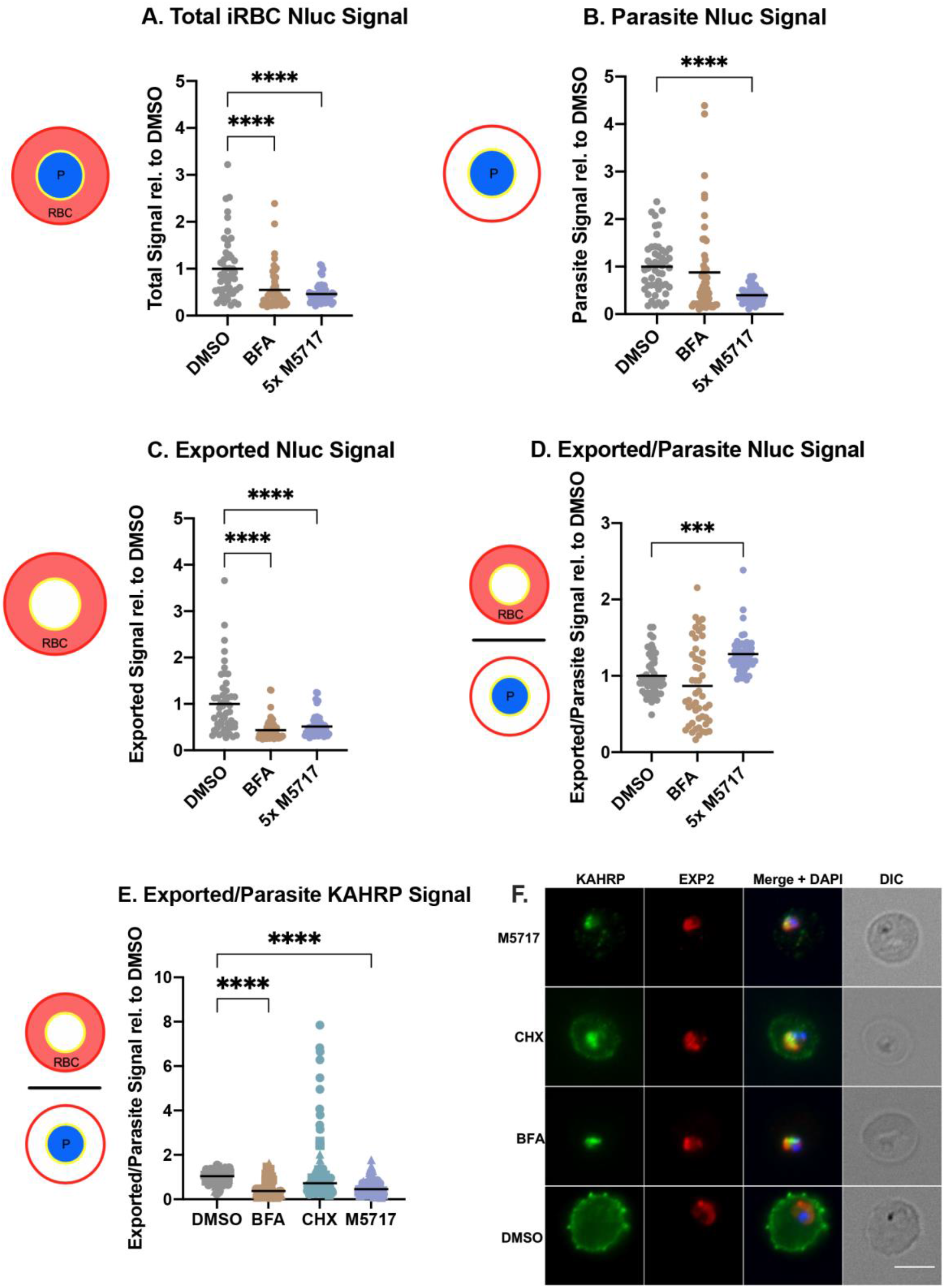
M5717 treated parasites have an overall reduced fluorescence intensity of exported proteins. Immunofluorescence microscopy was performed on Hyp1-Nluc parasites treated with M5717 (2.8 nM), BFA, (5 µg/mL) and DMSO (0.1%) for 5 h from 18 hpi (±2 h) (A-D); and on 3D7 parasites that had been treated similarly, with the addition of cycloheximide (100 µM, CHX) for 5 hours from 15 hpi (±1 h) (E-F). **(A-D)** Cells were probed with anti-Nluc IgG for exported Hyp1 and anti-EXP2 to mark the PVM. Anti-Nluc fluorescent signal was measured in the **(A)** whole iRBC, **(B)** the parasite and **(C)** the iRBC minus the signal in the parasite to show the amount of exported Nluc. **(D)** The ratio of exported/retained Nluc signal was thus determined. **(E)** The ratio of exported/retained KAHRP signal is determined from probing the cells with anti-KAHRP for exported signal and anti-EXP2 to mark the PVM. **(F)** Representative images of 3D7 parasites after 5 h of treatment, probed with anti-KAHRP (green) and anti-EXP2 (red) and DAPI (blue) to stain the nucleus. Treatment with M5717 at older stages results in higher signal in the RBC than the parasite vacuole, whilst younger parasites have reduced signal. Average signal was determined as the total fluorescence from each cell over the region from which it was measured. **(A-D)** N = 50 cells per treatment, **(E)** N = 3 biological replicates (represented by circle, triangle, and square symbols). Medians are presented as a black bar and an ordinary one-way ANOVA with multiple comparisons was performed with Prism software. ** p = 0.0011, *** p = 0.0005 and **** p = <0.0001. Scale bar = 5 µm.

To further investigate the effect of parasite age on the inhibition of protein synthesis and export, as well as examine the impact on cytoadherence capabilities of the parasites, younger 3D7 parasites at 15 hpi (± 1 h) were similarly treated for 5 hours, with the additional control of known translation inhibitor CHX. The cells were probed for EXP2 to mark the PVM again, and for knob associated histidine rich protein (KAHRP) as a component of the virulence complex involved in the mechanism of cytoadherence of iRBCs to the microvasculature (Taylor et al., 1987). Overall, the results supported the previous findings of fluorescence signal reduction in M5717 treated Hyp1-Nluc parasites (Figures 4, S4, S5). As predicted, when the parasites were treated from an age preceding export, the ratio of exported KAHRP signal to parasite signal was reduced, as opposed to the results seen in later stage parasites probed with Nluc (Figure 4D, E). This can be visualised in the representative IFA images where there is a noticeable reduction in the KAHRP signal (green) in the exported RBC compartment of parasites treated with M5717, as well as the translation inhibitor CHX (Figure 4F).

### 3.6 M5717 treatment has differential effects on the biomechanical properties of infected erythrocytes

New Permeability Pathways (NPPs) are formed in the iRBC membrane as the parasite develops into a trophozoite, allowing the uptake of nutrients from the plasma for the parasite’s use. NPPs are thought to be formed by the RhopH protein complex, following the export of these proteins from the parasite to the iRBC membrane (Nguitragool et al., 2011;Counihan et al., 2017;Ito et al., 2017;Sherling et al., 2017). If export of the RhopH proteins and formation of the RhopH complex at the iRBC membrane is inhibited by M5717, this could reduce the iRBC’s immunological profile, allowing these parasites to avoid clearance from the bloodstream (Shao et al., 2022). As iRBCs persisted in infected volunteers treated with M5717 we therefore measured the kinetics of NPP activity using a sorbitol lysis assay. Sorbitol enters the iRBC via the NPPs, thereby increasing the osmotic pressure within the RBC and causing hypotonic lysis (Figure 5A, left) (Wagner et al., 2003). If parasite proteins are not expressed, and thus fail to reach the plasma membrane NPPs would not be formed and lysis would not occur (Counihan et al., 2017). To measure iRBC lysis, Hyp1-Nluc parasites were used as they release bioluminescent protein upon RBC lysis. Synchronous 22-hour parasites (± 2 h) were treated for 5 hours with 5x EC_50_ M5717, a known NPP inhibitor furosemide (Gilson et al., 2019) or a DMSO vehicle control. The cultures were either lysed with Nano-Glo containing sorbitol lysis buffer or phosphate buffered saline (PBS) as a control for the background signal. Bioluminescence was measured every 3 minutes over 43.5 minutes to determine the NPP kinetics by fitting the data with an exponential growth equation. This resulted in a constant rate (k) value that was normalised by removing background signal and comparing against the DMSO control. Furosemide showed potent inhibition of the NPP activity (Figure 5A, right). Whilst reduced signal was observed after M5717 exposure, the difference was not significant and more likely reflects the overall effect of M5717 on protein synthesis and not direct inhibition of the NPPs (Figure 5A).

**Figure 5.**
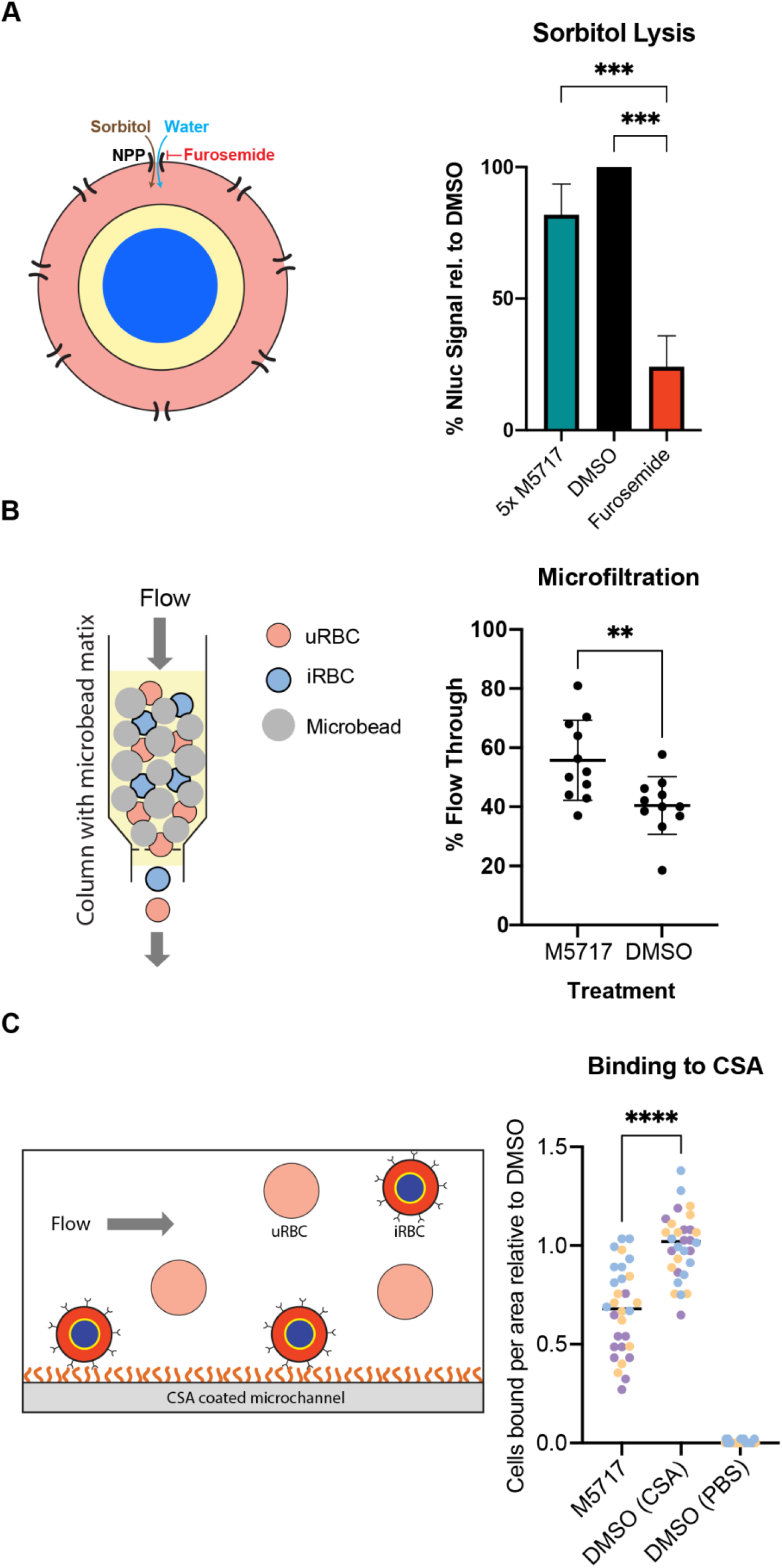
Effect of M5717 on parasite induced biophysical properties of infected RBCs. **(A, Left)**. Diagram of parasite iRBC showing how the movement of isotonic sorbitol followed by water into the iRBC via the new permeability pathway (NPP) causes the iRBCs to swell and rupture. **(Right)** iRBCs (22 hpi, ±2 h) were treated for 5 h with either 5x EC_50_ M5717 (2.8 nM), NPP inhibitor furosemide (100 µM) or 0.1% DMSO. The cultures were treated with isotonic sorbitol buffer containing Nano-Glo substrate or control 1x PBS. Bioluminescence was measured every three minutes to determine the kinetics of sorbitol induced lysis with DMSO normalised to 100%. Known NPP inhibitor furosemide strongly inhibits NPP activity while M5717 does not. Error bars represent standard deviation of three biological replicates with three technical replicates each. An ordinary one-way ANOVA with multiple comparisons was performed with Prism software, *** p = 0.0008. **(B, left)** Diagram showing how iRBCs can be pumped through a bed of spleen-mimicking microbeads with passage of the stiffer iRBCs being impeded more than the flexible iRBCs and uRBCs. **(Right)** Transgenic MSP1-Nluc (15 hpi, ±1 h) iRBCs were treated with 5x EC_50_ M5717 (2.8 nM) or vehicle control DMSO (<0.001%) for 5 h. The percentage of parasites that flowed through the beads were calculated by comparing the parasitemia of input samples to those that passed through. M5717 treatment allowed a higher proportion of iRBCs to flow through than the DMSO control, indicating M5717 inhibited the capacity of the parasites to stiffen the iRBCs. Error bars represent standard deviation of three biological replicates, where N = 1000 cells counted. A Welch’s t test was performed with Prism software, ** p = 0.0072. **(C, left)** Diagram showing CS2 parasites presenting Var2CSA at the iRBC surface flowing over and binding to a CSA-coated surface. **(Right)** Synchronous iRBCs (15-20 hpi) were treated for 5 h with either 5x EC_50_ M5717 (2.8 nM) or DMSO (<0.001%) and were flowed across CSA (100 µg/mL) coated microchannel slides to allow substrate binding to occur. A control was performed where DMSO treated parasites were flowed across slides treated with 1x PBS lacking CSA to demonstrate binding was specific for CSA. To quantitate binding, adherent cells were counted in 10 random fields for 3 biological repeats of each condition, and then normalised against the DMSO (CSA). Treatment with M5717 resulted in less cells successfully adhering to the CSA coated slides. Medians are presented as a black bar and an ordinary one-way ANOVA with multiple comparisons was performed with Prism software, **** p = <0.0001.

Exported parasite proteins interact with the RBC cytoskeleton, altering its structure and in turn increase the rigidity of iRBCs compared to highly flexible uninfected RBCs (Glenister et al., 2002). The inhibition of protein synthesis caused by M5717 could reduce the capacity of parasites to make their iRBCs more rigid so that they pass uncleared through the spleen, thereby remaining in circulation. To test this, a second parasite line that secretes Nluc into its parasitophorous vacuole (MSP1-Nluc) was used in an iRBC rigidity assay (Deplaine et al., 2011;Looker et al., 2022). Here 15 hpi (± 1 h) iRBCs that had been treated with compounds were flowed through a bed of metallic microbeads through which only flexible cells can pass (Figure 5B, left). Prior to passage through this ‘spleen in a tube’ the parasites were treated for 5 hours with 5x EC_50_ M5717 or DMSO. The proportion of iRBCs relative to uninfected RBCs that passed through the microbeads was counted by light microscopy. Results showed that M5717 treatment resulted in relatively more iRBCs passing through the matrix, confirming that the compound slows stiffening of iRBC (Figure 5B, right).

Survival of *P. falciparum* iRBCs in the circulation is linked to their ability to avoid splenic clearance by sequestering in the deep vasculature. This is mediated by the formation of specialised structures beneath the RBC plasma membrane, termed knobs, which mediate specific receptor-ligand interactions with vascular endothelial cells. Knob formation is mediated by trafficking of exported parasite proteins to the RBC membrane (Magowan et al., 1988). As previously shown, the synthesis and export of KAHRP, a knob-associated protein, is reduced in M5717 treated parasites (Figures 4, S5). To investigate further if M5717 inhibits cytoadhesion, we used CS2, a line of *P. falciparum* parasites that had been selected for cytoadherence to the endothelial receptor chondroitin sulphate-A (CSA), an adhesin which is associated with placental malaria (Clausen et al., 2012). CS2 parasites at 15-20 hpi were treated for 5 hours with either 5x EC_50_ M5717 or DMSO before being flowed across CSA coated microchannel slides (Figure 5C, left). To quantify cytoadherence, the number of iRBCs bound to the slide were counted in 10 random fields for 3 biological repeats of each condition. Treatment with M5717 resulted in fewer cells adhering to the CSA-coated slides compared to the DMSO control, per field (281 x 178 µm) (Figure 5C, right, Figure S6). To ensure binding to CSA was specific, a control was performed where DMSO treated parasites were flowed across slides not treated with PBS containing no CSA where there was negligible binding (Figure 5C).

## 4 Discussion

Our results indicate persistence of iRBCs in the blood stream following M5717 administration can be explained by the drug preventing parasites from deploying RBC-modifying proteins that would increase RBC rigidity thereby reducing splenic clearance, while simultaneously inhibiting sequestration, ultimately allowing persistence in the circulation. This effect of M5717 appears to be a result of direct inhibition of the synthesis of exported proteins, rather than by specifically blocking protein export to the iRBC. This is supported by the bioluminescent export assay and fluorescence microscopy results.

In seeking to understand how the inhibition of synthesis of exported proteins could facilitate the persistence of iRBCs in the blood stream we examined three biophysical properties of iRBCs known to be caused by exported proteins namely, plasma nutrient uptake, the rigidity and cytoadherent properties of iRBCs. The installation of the RhopH/NPP complex at the plasma membrane of the iRBC did not appear to significantly reduce following M5717 treatment, using a nanoluciferase-based sorbitol cell lysis assay. This is probably because RhopH2 and RhopH3 are largely synthesised during merozoite development, and are rapidly transported into newly infected RBCs shortly after export, well before we performed treatment with M5717 (Pasternak and Dzikowski, 2009). M5717 treatment did not commence until 22 hours after invasion, by which time the RhopH complex had probably already been formed. We note that PTEX is expressed similarly as the RhopH proteins and a functional translocon was likely in place soon after invasion (de Koning-Ward et al., 2009;Bullen et al., 2012).

Some exported proteins such as KAHRP are known to contribute to knob formation and increase the stiffness of iRBC (Pei et al., 2005;Rug et al., 2006). Therefore, we treated mid-ring infected iRBCs with M5717 at a time in their lifecycle when they were still synthesising KAHRP and other cytoskeleton-binding proteins, to observe by immunofluorescence any changes in signal intensity. We found an overall reduction in KAHRP signal after treatment with M5717, resulting in lower levels of KAHRP export to the RBC membrane. To further determine if this reduced iRBC stiffness and to determine if M5717 treatment ultimately caused the iRBCs to become less rigid than the inactive control treatment, we performed filtration through a microbead matrix. In an *in vivo* setting, this could prevent splenic clearance of iRBCs containing younger trophozoites, thereby permitting them to continue circulating (Deplaine et al., 2011). Flexible iRBCs would cytoadhere less, reducing their capacity to sequester in deep tissues, thus allowing them to remain in circulation. Older, larger trophozoites and schizonts would probably still be rigid enough to be trapped by the spleen.

Finally, we attempted to directly measure the cytoadherence capacity of M5717 treated iRBCs under flow conditions by microscopy. In this experiment we used CS2 parasites that have been shown to bind well to CSA-coated microcapillaries (Cooke et al., 1996;Looker et al., 2022), and observed that cytoadherence of M517 treated iRBCs was inhibited. Whether M5717 directly reduced the production of the Var2CSA variant PfEMP1, reduced the formation of cytoadherence surface knobs on the iRBC surface from which Var2CSA is presented or simply reduced the rigidity of the whole iRBC thereby decreasing cytoadherence was not determined. In conclusion, the persistence of iRBCs in the blood stream after M5171 administration is likely due to the inability of M51717 treated parasites to produce the exported proteins necessary to alter the biophysical properties of iRBCs. Study of the in vivo effect of slowed clearance of parasites on the clinical effects of severe malaria in the setting of high parasitemia, and possible changes to the immune response to infection following treatment with this drug are worth investigation.

## Supporting information

All supplementary figures

## Data availability statement

The original contributions presented in the study are included in the article/Supplementary Material. Further inquiries can be directed to the corresponding author.

## Author contributions

MPS, OL, MR, JM and PRG conceived this research and designed the experiments. MPS, OL, MR and DKK conducted the experiments and generated the figures. MPS, OL, MR, JM and PRG wrote the manuscript. MPS, OL, MWAD, JM and PRG edited the manuscript. BSC, PRG, CO and JM provided funding. All authors have approved the submitted manuscript.

## Acknowledgements

We are grateful to the Australian Red Cross Lifeblood for the provision of human blood and thankful for funding from the Victorian Operational Infrastructure Support Program received by the Burnet Institute. We thank Merck KGaA, Darmstadt, Germany for M5717 compound. We would like to thank the Burnet Cell Imaging Facility for assistance and access to equipment.

