## Supplementary material for "The delayed bloodstream clearance of *Plasmodium falciparum* parasites after M5717 treatment is attributable to the inability to modify their red blood cell hosts": All supplementary figures

**SUPPLEMENTARY DATA FILE**

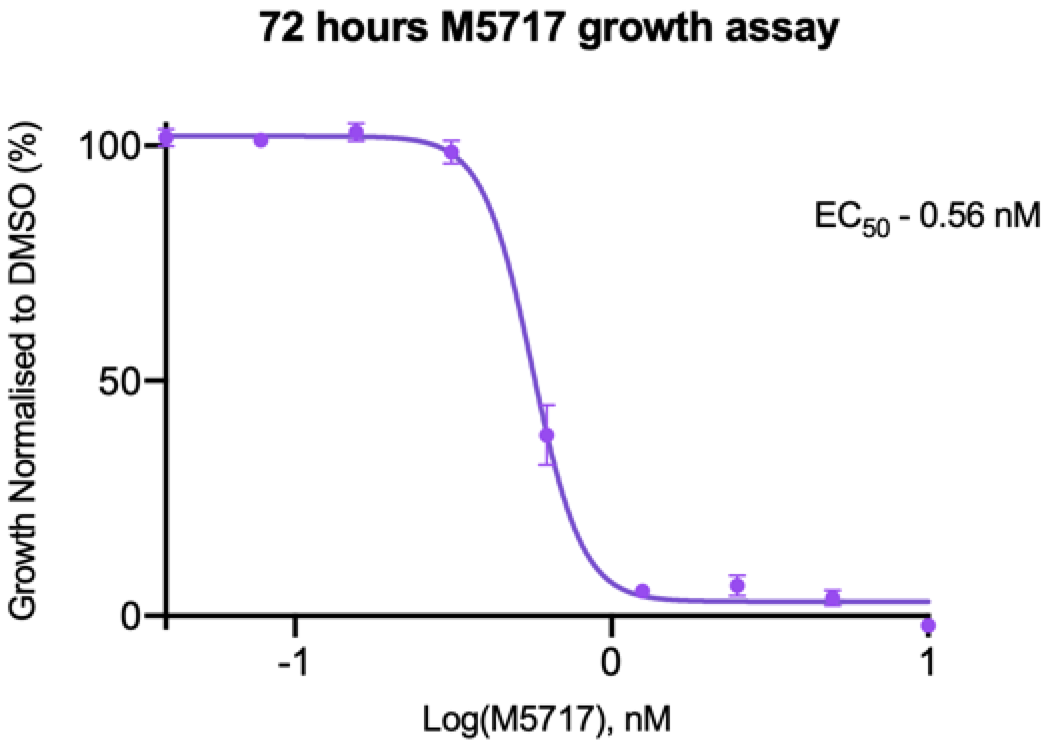

**Supplementary Figure 1** Growth inhibition assay of M5717 on *P. falciparum* 3D7 parasites indicated the $EC_{50}$ was 0.56 nM. A 72-hour growth assay was initiated at ring stage with a 2-fold, 9-point dilution series of M5717, starting at 10 nM. Controls were performed using uninfected RBCs and DMSO vehicle. Parasite viability was determined by measurement of lactate dehydrogenase activity. The $EC_{50}$ was calculated using Graphpad Prism software. Error bars represent standard deviation of three technical replicates.

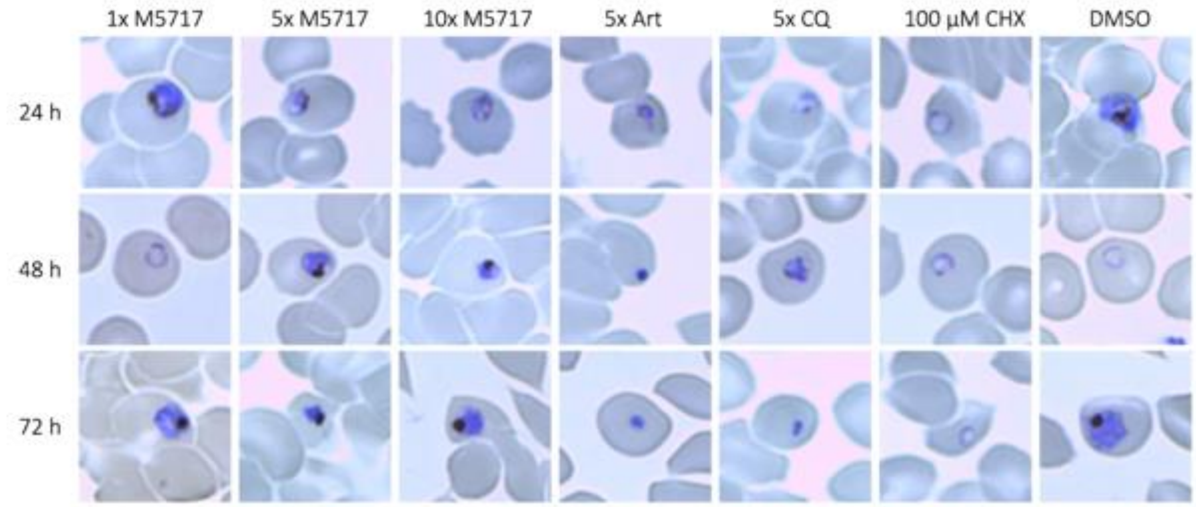

**Supplementary Figure 2** Giemsa-stained thin blood smears of *P. falciparum* Hyp1-Nluc ring stage parasites treated with M5717 persist for 72 h with a trophozoite-like appearance. To observe the viability and morphological effect of M5717 on *P. falciparum* Hyp1-Nluc synchronous ring stage parasites, they were treated separately with either 1x, 5x or 10x $EC_{50}$ of M5717 (0.56 nM, 2.8 nM or 5.6 nM). Parasites were also treated with the known antimalarials artemisinin (Art, 40 nM), chloroquine (CQ, 64.5 nM), translation inhibitor cycloheximide (CHX), and the vehicle control DMSO (0.05%). Giemsa-stained thin blood smears after 24, 48 and 72 h treatment are shown. After 24 h growth delays became visible in Art and CQ and at higher M5717 concentrations. In contrast, 1x M5717 and DMSO progressed through invasion to from new rings. By 72 h, the Art and CQ parasites became pyknotic and the M5717 appeared to have stalled as abnormal trophozoites

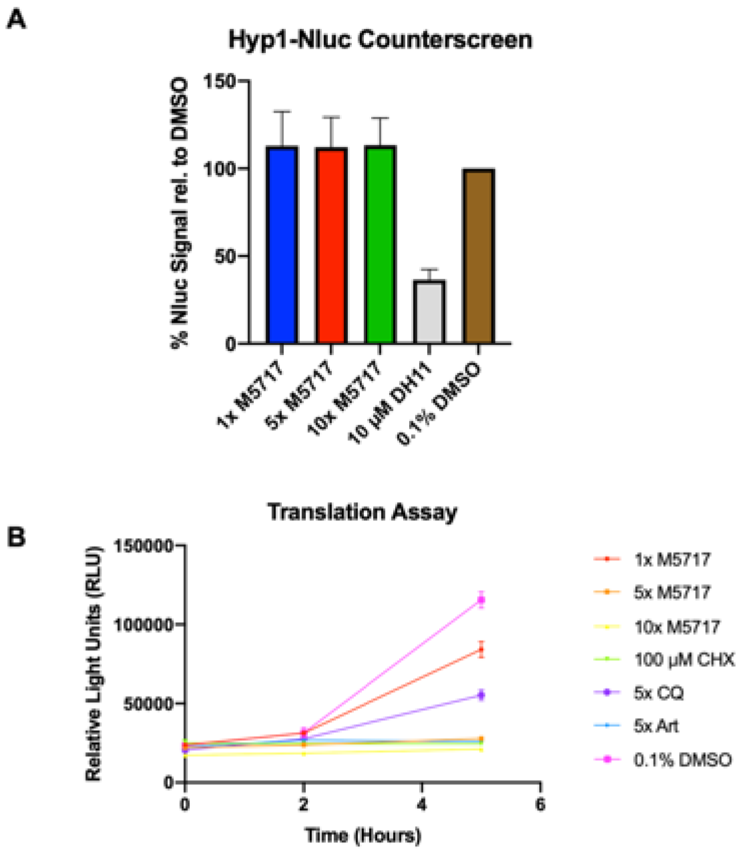

**Supplementary Figure 3** M5717 does not directly inhibit nanoluciferase activity but does inhibit translation of the enzyme. **(A)** Bioluminescence of the Hyp1-Nluc reporter was not inhibited by M5717 at 1x, 5x or 10x M5717 $EC_{50}$ (0.56 nM, 2.8 nM, 5.6 nM) or a DMSO vehicle control (0.1%) normalised to 100%. Medicines for Malaria Box compound MMV665972 (DH11) is a known inhibitor of nanoluciferase. Results represent the average of three technical replicates, error bars show standard deviation. **(B)** M5717 suppresses Hyp1-Nluc expression in a concentration dependent manner in young trophozoite stage parasites. Synchronous *P. falciparum* Hyp1-Nluc (18-20 hpi) parasite iRBCs were treated with three concentrations of M5717 (1x $EC_{50}$, 5x $EC_{50}$, 10x $EC_{50}$), artemisinin (Art, 40 nM), chloroquine (CQ, 64.5 nM), cycloheximide (CHX, 100 µM) or vehicle control DMSO (0.1%). Samples were collected after 2 and 5 h of treatment and nanoluciferase bioluminescence was measured in relative light units (RLU) after the addition of Nano-Glo. Error bars indicate the standard deviation of three technical replicates.

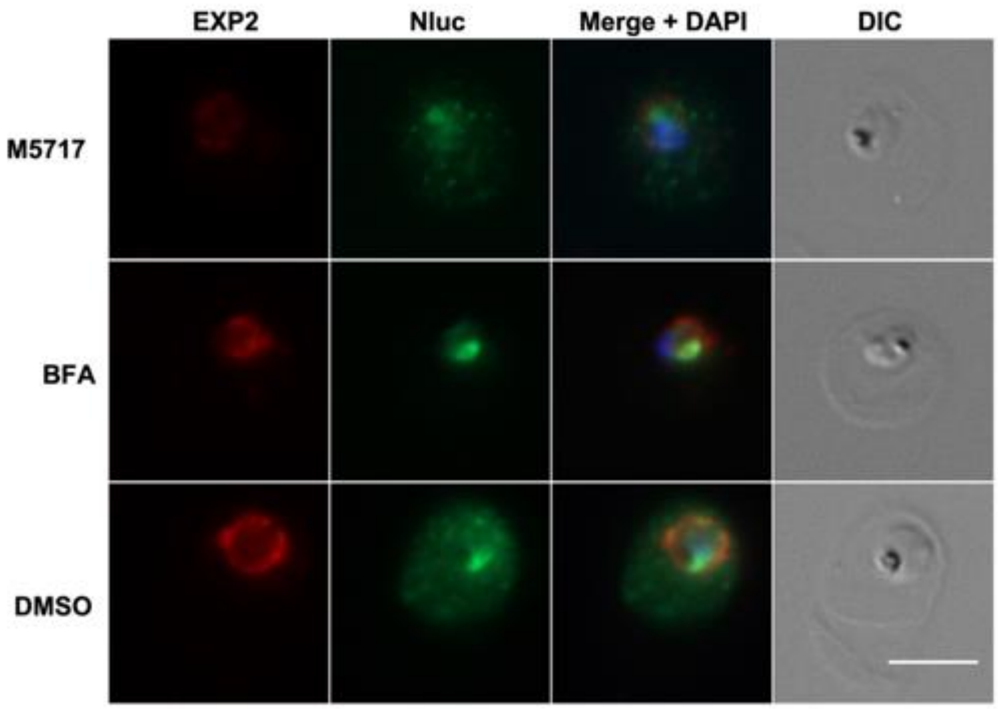

**Supplementary Figure 4** Representative IFA images of treated Hyp1-Nluc parasites. An immunofluorescence assay was performed on Hyp1-Nluc parasites at 18 hpi (±2 h) treated with M5717 (2.8 nM), the export inhibitor brefeldin A (BFA, 5 µg/mL) and vehicle control DMSO (0.1%) for 5 h. The cells were probed with anti-Nluc (green) for exported Hyp1 and anti-EXP2 (red) to mark the PVM. Signal intensity is reduced in BFA and M5717 treated parasites and retained within the PV by the export inhibitor BFA. Scale bar represents 5 µm.

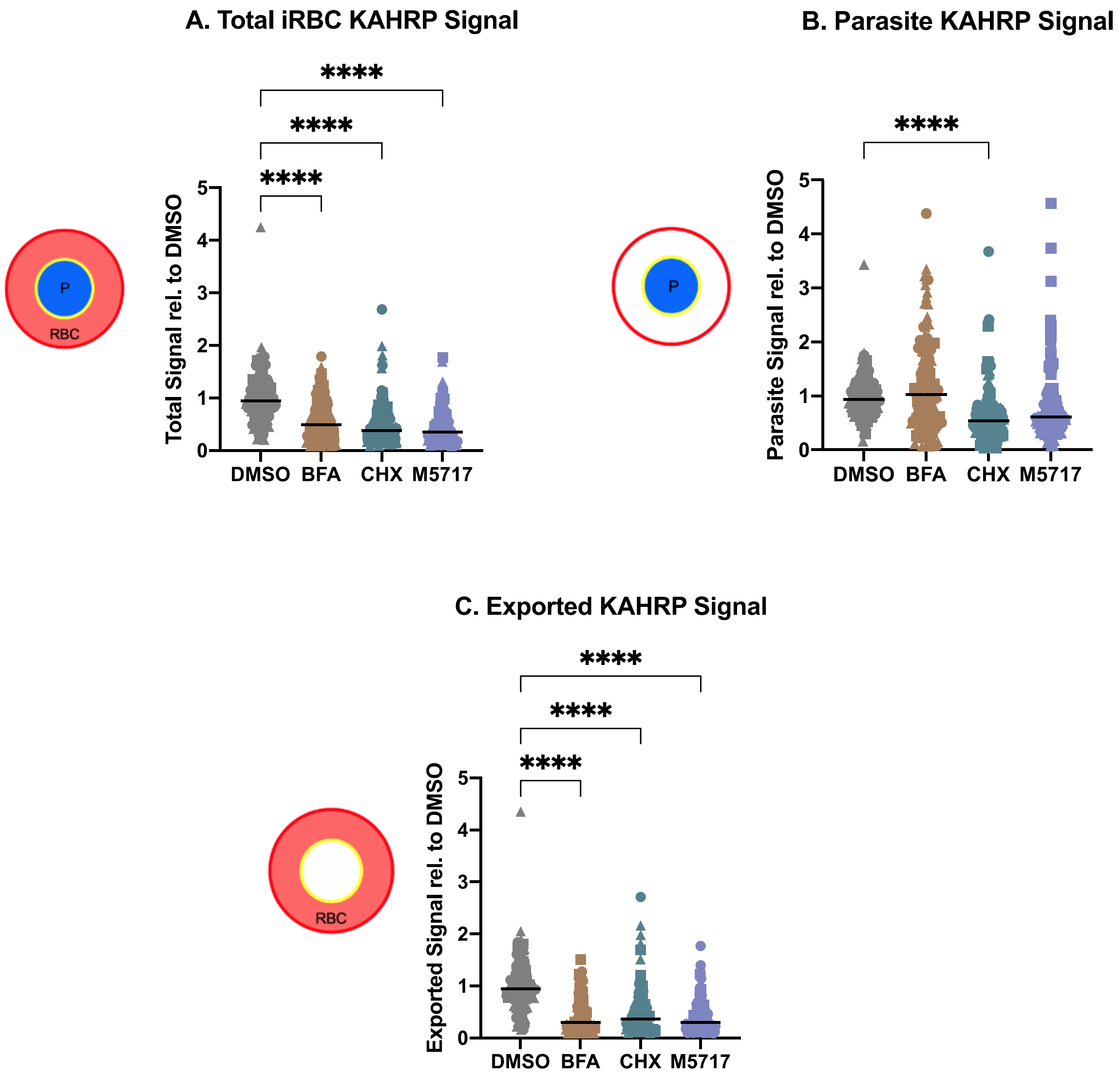

**Supplementary Figure 5** M5717 treatment reduces KAHRP fluorescence intensity. Immunofluorescence microscopy was performed on 3D7 parasites treated with M5717 (2.8 nM), BFA, (5 µg/mL), cycloheximide (100 µM, CHX) and DMSO (0.1%) for 5 h from 15 hpi (±1 h). Cells were probed with anti-KAHRP and anti-EXP2 to mark the PVM. (A) Anti-KAHRP fluorescence was measured in the **(A)** whole iRBC, **(B)** the parasite and **(C)** the iRBC minus the signal in the parasite to show the amount of exported protein. Treatment with M5717 results in decreased anti-KAHRP signal overall. Average signal was determined as the total fluorescence from each cell over the region from which it was measured. N = 3 biological replicates (represented by circle, triangle and square symbols). The Median is presented as a black bar and an ordinary one-way ANOVA with multiple comparisons was performed with Prism software. **** p = <0.0001.

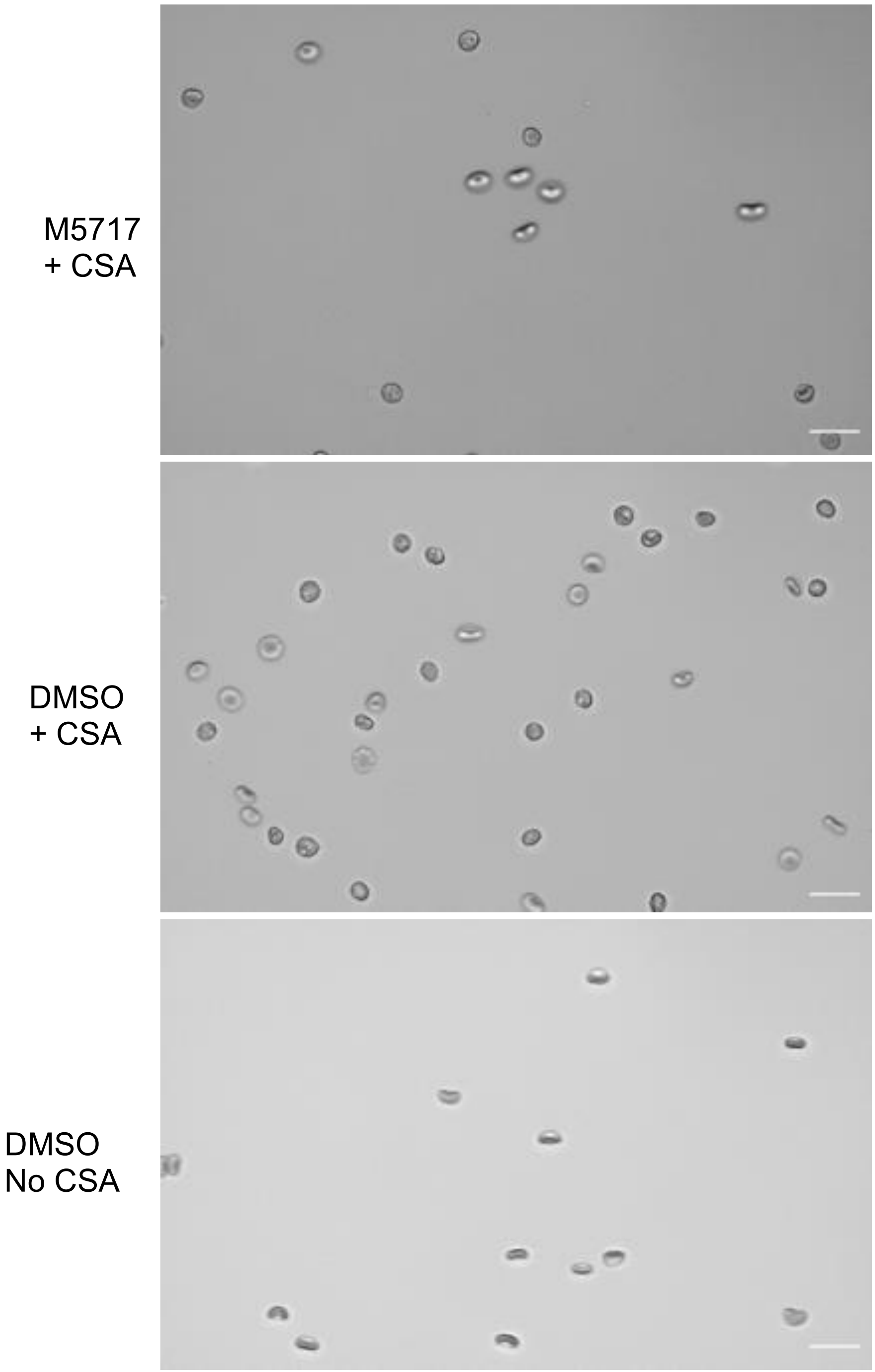

**Supplementary Figure 6** Representative fields of view for microchannel cytoadherance experiment. Microscopy images showing how CS2 parasite iRBCs can export Var2CSA onto the iRBC surface to facilitate their cytoadherence to CSA-coated substrate. Parasite iRBCs (15-20 hpi) were treated for 5 h with either M5717 (2.8 nM) or DMSO (<0.001%) and were flowed across CSA (100 µg/mL) in PBS coated microchannel slides to allow substrate binding to occur. A control was performed where DMSO treated parasites were flowed across slides treated with PBS only (13) (Bottom) to demonstrate binding was specific for CSA. To quantitate binding, adherent cells were counted in 10 random fields for 3 biological repeats of each condition, and then normalised against the DMSO. Treatment with M5717 resulted in less cells successfully adhering to the CSA coated slides. The scale bar represents 20 µm.
